# Breakdown of gametophytic self-incompatibility in subdivided populations

**DOI:** 10.1101/444125

**Authors:** Thomas Brom, Vincent Castric, Sylvain Billiard

**Affiliations:** Univ. Lille, UMR 8198 - Evo-Eco-Paleo, F-59000 Lille, France; CNRS, UMR 8198, F-59000 Lille, France

**Keywords:** mating system, inbreeding depression, dispersal, fragmentation

## Abstract

Many hermaphroditic flowering plants species possess a genetic self-incompatibility (SI) system that prevents self-fertilization and is typically controlled by a single multiallelic locus, the S-locus. The conditions under which SI can be stably maintained in single isolated populations are well known and depend chiefly on the level of inbreeding depression and the number of SI alleles segregating at the S-locus. However, while both the number of SI alleles and the level of inbreeding depression are potentially affected by population subdivision, the conditions for the maintenance of SI in subdivided populations remain to be studied. In this paper, we combine analytical predictions and two different individual-based simulation models to show that population subdivision can severely compromise the maintenance of SI. Under the conditions we explored, this effect is mainly driven by the decrease of the local diversity of SI alleles rather than by a change in the dynamics of inbreeding depression. We discuss the implications of our results for the interpretation of empirical data on the loss of SI in natural populations.

**Data accessibility statement:** No data to be archived

## Introduction

Around half of hermaphroditic flowering plants species have the capacity to prevent selfing and enforce outcrossing through a self-incompatibility (SI) system (Ivanov et al. 2010; Gibbs 2014). SI systems have evolved multiple times with different molecular mechanisms (Steinbachs and Holsinger 2002; Castric and Vekemans 2004). In many species, SI specificities are controlled by a large allelic series at a single multiallelic locus, the S-locus (Takayama and Isogai 2005), and SI is actively achieved by a rejection reaction that prevents self-fertilization as well as fertilization between individuals expressing cognate specificities (De Nettancourt 1997; Gibbs 2014).

Transitions from outcrossing to selfing are widespread in the flowering plants (Stebbins 1974; Barrett 2002; Igic et al. 2008; Shimizu and Tsuchimatsu 2015) and there has been considerable theoretical interest to understand the conditions under which self-compatible (SC) mutants can invade a SI population, a prerequisite for the evolution of selfing. Briefly, the conditions for maintenance of SI are expected to reflect the balance between its advantages (mainly the prevention of the deleterious effects of inbreeding depression), and the variety of its costs (including the cost of outcrossing Fisher (1941) and the fact that SI acts as a barrier to fertilization between individuals expressing cognate specificities, and as such can impair mating success). In a seminal study, Charlesworth and Charlesworth (1979) showed that SC mutants are expected to invade a SI population when the intensity of inbreeding depression is below some threshold. In addition, they showed that the invasion was more straightforward when the number of segregating SI alleles is limited, because SI pollen then suffers from being rejected by individuals with the same SI allele while the SC mutants have universal compatibility and therefore enjoy a comparative advantage through the male reproductive function. More recently, as the segregation of SC mutants may be expected to allow purging of the genetic load (Glémin 2003) and may thus decrease inbreeding depression, Gervais et al. (2014) explicitly modeled the accumulation of partially recessive mildly deleterious mutations, thus considering inbreeding depression as a dynamic variable rather than a fixed parameter. They showed that allowing inbreeding depression to vary had only a marginal impact on the qualitative pattern beyond a slight decrease of the parameter range under which SI is maintained.

Subdivision is one of the most common features of natural populations (Selander 1970; Loveless and Hamrick 1984; Slatkin 1987), but the breakdown of SI in a spatially structured population has never been theoretically investigated. On the one hand, a general effect of the strong balancing selection acting on the SI genes (Wright 1939) is to maintain among-demes differentiation at a low level. Thus, SI alleles should be distributed more homogeneously among local demes than variants at the genomic background (Schierup 1998) and the effect of population subdivision could be negligible. On the other hand, there are several reasons to believe that the dynamics of SI and SC mutants may differ from that in a single isolated population. First, population subdivision should have an effect on the diversity of SI alleles maintained, both locally and globally (Schierup 1998). The effect of population subdivision on the global and local diversity of SI alleles is however not trivial. Indeed, while local diversity always decreases with increasing population subdivision, Schierup (1998) showed that the global diversity of SI alleles does not vary monotonously with pollen dispersal. In fact, strong subdivision may allow different sets of SI alleles to stably segregate in the different subpopulations, such that there are conditions where intermediate levels of isolation are associated with the lowest global diversity of SI alleles. Thus, the cross-compatibility advantage for pollen of a SC mutant can be expected to depend on a combination of pollen dispersal rate and the local and global diversity of SI alleles whose interaction is potentially complex and has not been explored yet. Second, population subdivision adds complexity to the dynamics of inbreeding depression, because in subdivided populations the decline in fitness experienced by a selfer depends on the relative fitnesses of outcrossed individuals within the demes and outcrossed individuals between the demes (Glémin et al. 2003). An increasing subdivision has an adverse effect on within-demes and between-demes inbreeding depressions, decreasing the first and increasing the second in most cases (Theodorou and Couvet 2002; Glémin et al. 2003; Roze and Rousset 2004). Furthermore, population structure should affect the impact of selfing on the purging process of the mutation load. In subdivided populations, self-fertilization always decreases within-demes inbreeding depression relative to random mating whereas high and low self-fertilization rates increase and decrease respectively between-demes inbreeding depression (Roze and Rousset 2004). Finally, when inbreeding depression depends on multiple loci, selective interference can reduce the effect of spatial structure and increases within-deme inbreeding depression (Roze 2015). Overall, due to the multiple processes cited above, it is not straightforward to predict the dynamics of inbreeding depression in a subdivided population when SC mutants are introduced.

In this study, we determined the conditions under which a subdivided population with gametophytic SI can resist invasion by a SC mutant. We combined analytical predictions and individual-based simulation models to compare the conditions under which SI can be maintained with different levels of subdivision. We explored the impact of the mechanisms mediating this effect, namely the diversity of SI alleles and the dynamics of inbreeding depression.

## Methods

We assumed a population of a hermaphroditic plant species with gametophytic SI (GSI). The population was subdivided in equally-sized demes and dispersal occurred with equal probability between each pair of demes, following an island model. We first performed an analytical analysis and then built two individual-based models: one with constant inbreeding depression and one with explicit modelling of deleterious mutations to allow inbreeding depression to vary dynamically. The analytical model allowed us to specifically investigate the effect of pollen flow on competition among pollen expressing the different SI and SC alleles, independently of its effect on gene flow *i.e.* on the number of SI alleles. Comparison between the results of the models with constant *vs.* variable inbreeding depression allowed us to investigate the importance of the purging process of deleterious mutations.

### Analytical model

We adapted the model of Gervais et al. (2014) by assuming a metapopulation of an annual plant made of an infinite number of infinitely large demes interconnected by pollen flow. We focused on one focal deme of this metapopulation and made the following assumptions: (i) the number of SI alleles *n* in the focal deme is constant and all S-alleles have equal frequencies; (ii) pollen disperse at rate *d_p_*, which we define as the proportion of outcross pollen each plant of the focal population sends away in the metapopulation and the proportion of (outcross) pollen the metapopulation sends to the focal deme; (iii) the effect of pollen flow from the focal deme towards the metapopulation is neglected, *i.e.* a SC allele cannot invade the metapopulation and pollen from the metapopulation only contains SI alleles; (iv) the number of SI alleles in the metapopulation is much larger than in the focal deme such that any immigrant pollen is compatible with all individuals of the focal deme.

To investigate the fate of SC mutants introduced at low frequency in the focal population of SI individuals, we followed the frequency *x_i_* of four types of individuals that differ by S-locus genotype and by origin. We denote *x*_1_ and *x*_2_ the frequency of individuals with two SC alleles (SC/SC), produced by selfing and by outcrossing respectively; *x*_3_ and *x*_4_ the frequency of individuals with a SC and a SI allele (SC/SI), produced by selfing and by outcrossing respectively. It follows that the frequency in the focal deme of individuals with two self-incompatible alleles (SI/SI) is 1 − *x*_1_ − *x*_2_ − *x*_3_ − *x*_4_. To determine the male and female gamete production *W_i_* of each type of individuals *i*, we assumed that there is no heterosis, *i.e.* outcrossed individuals with a father from the focal population and outcrossed individuals with a father from the metapopulation have an identical gamete production *W_o_*, defined as the gamete production of outcrossed individuals. Thus *W*_2_ = *W*_4_ = *W_o_* = 1, with *W*_2_ and *W*_4_ gamete production of outcrossed individuals, with two and one SC alleles respectively. Selfed individuals with two and one SC alleles have gamete production *W*_1_ and *W*_3_ respectively. These latter individuals suffer from inbreeding depression *δ*, assumed to be constant and acting on gamete production only. Thus *W*_1_ = *W*_3_ = *W_s_* = 1 − *δ* with *W_s_* the gamete production of selfed individuals. The mean gamete production in the focal deme is thus *W̄* = *W _s_* (*x*_1_+ *x*_3_)+*W_o_* (1*− x*_1_ *− x*_3_) = 1 − *δ* (*x*_1_+ *x*_3_). The frequency *q* of SC alleles in male gametes is

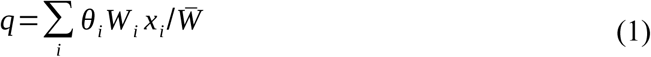

and the frequency of each SI allele in the male gametes is

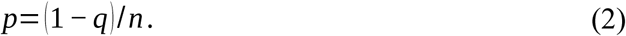

For each type of individuals, we determined the selfing, within-deme outcrossing and between-demes outcrossing rates, which we define as the proportion of female gametes fertilized by self-pollen, by pollen from the focal deme or by pollen from the metapopulation, respectively. These quantities depend on the amount of compatible pollen received from three sources: i) self-pollen: *α θ_i_ W_i_*, ii) outcross pollen from individuals from the focal deme: (1 −*α*)(1− *d p*)*W̄* (1 −2 *p*)≃ (1 − *α*) (1− *d p*)*W̄* under the assumption that the number of SI alleles is large and iii) outcross pollen from individuals from the rest of the metapopulation: (1 −*α*)*d_p_ W_o_*, where *α* is the proportion of self-pollen and *θ_i_* the proportion of self-compatible self-pollen for each type of individuals (*θ*_1_ = *θ*_2_ = 1 and *θ*_3_ = *θ*_4_ = 1/ 2). Then, the effective selfing rate of individuals *i* of frequency *x_i_* is

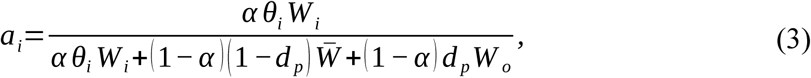

its effective within-deme outcrossing rate is

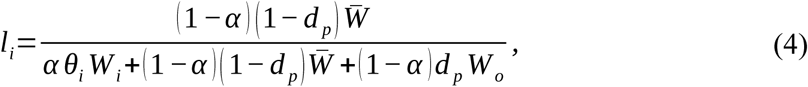

and its between-demes outcrossing rate is

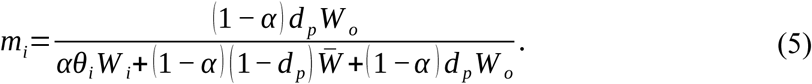

It follows that the frequencies at the next generation of the four types of individuals are given by

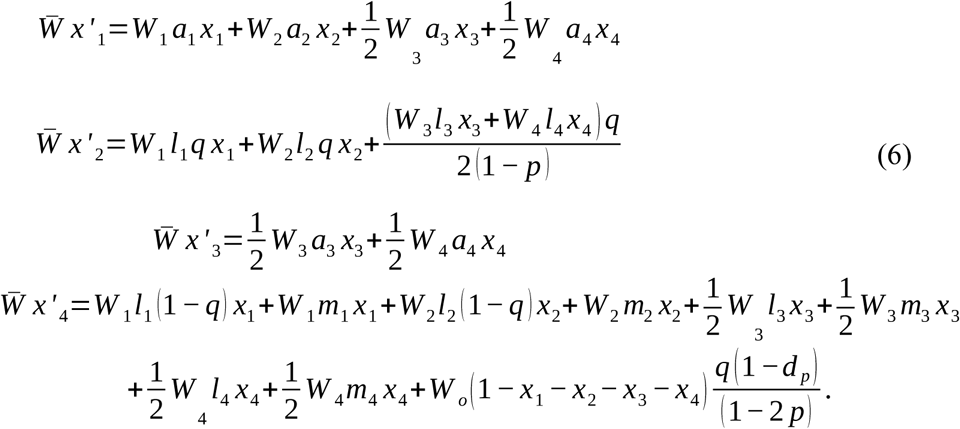

Note that with *d_p_* = 0 the equations above are exactly the same as in a panmictic population (Gervais et al. 2014).

Following Gervais et al. (2014), we used the equation of the frequencies of each type of individual at the next generation (Eq. 6) to perform a local stability analysis, whereby we determined for different *d_p_, α* and *n*, the values of *δ* for which the SC allele was expected to increase in frequency when rare. We simplified the system by considering *x*_2_ (the frequency of individuals with two SC alleles from outcrossing) negligible when SC is rare. By simplifying Eq. (6) to the first order in *x*_1_, *x*_3_ and *x*_4_ we get a stability matrix *A* such that (*x*_1_, *x*_3_, *x*_4_)’ = *A*. (*x*_1_, *x*_3_, *x*_4_). We used Routh-Hurwitz conditions to determine the stability of the equilibrium when SC is rare without having to calculate the eigenvalues of the stability matrix explicitly (Otto and Day 2007). For different values of *d_p_*, *α*, *n*, we solved numerically Routh-Hurwitz conditions, using Mathematica ver. 11.3.0.0 (Wolfram Research), to find the values of *δ* for which the equilibrium is unstable and the SC allele increases in frequency.

### Individual-based model

To complement the analytical model in more complex settings, we then ran two different sets of individual-based simulations, with either constant or dynamic inbreeding depression. We considered diploid hermaphroditic plants with non-overlapping generations in a metapopulation of constant size *N* divided in *D* equally-sized demes. During the first steps of the simulations, we introduced no SC mutant and all individuals were obligate outcrossers. At the beginning of the simulations, we randomly drew for each individual two different SI alleles among *S* different possible SI alleles. Individuals in the population were then formed each generation by random draws with replacement of gametes from two parents from the previous generation proportionally to their gamete production. The female gamete was drawn from the local deme and the male gamete (pollen) was either drawn from the same deme as that of the female gamete with probability 1 *−d_p_*, or from another deme with probability *d_p_* the pollen dispersal rate. Pollen were drawn repeatedly until a compatible pollen was found (no pollen limitation). Each inherited SI allele had a probability *U _SI_* to mutate to another SI allele among the *S* possible different SI alleles. Allowing this type of mutation provides the conditions for long-term maintenance of SI allele’s diversity against loss by genetic drift. Later in the simulations, we recurrently introduced SC alleles into the metapopulation with probability *U _SC_* for each SI allele. When an individual *i* from the local deme *k* was drawn as a female, it produced a selfed offspring with probability *a_i_* given by:

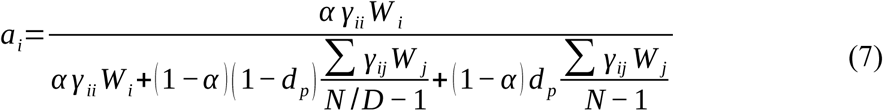

where *γ_ii_* is 0, 1 or 2, the number of SC alleles of individual *i*, *γ_ij_* the number of S-alleles of individual *j* compatible with the S-alleles of individual *i*, *W _i_* the pollen production of individual *i*, *W _j_* the pollen production of individual *j*. In case of selfing, if the selfing individual had only one SC allele, this SC allele was necessarily the paternal S-allele. If individuals had two SC alleles we randomly chose one of them. 1 −*a_i_* was then the outcrossing rate of individual *i* and was equal to 1 for individuals with no SC allele (*γ_ii_* = 0).

Note that when *d_p_* = 0, we recovered the same situation as in isolated populations (Gervais et al. 2014) for the same deme size. Note also that in contrast to the analytical model, the dispersed pollen enters in competition with other pollen in other demes. Comparing results between the analytical model and simulations will thus give insights about the role of SC pollen export, especially for high dispersal. Similar results between simulations and analytical model would suggest that the SC pollen flow between demes has a negligible effect.

#### Modeling of inbreeding depression

In the first model we considered constant inbreeding depression, and individuals from outcrossing had a relative gamete production of 1 whereas individuals from selfing (male and female gametes from a same individual) had a relative gamete production of 1 − *δ* with *δ* the constant inbreeding cost.

In the second model, we followed the framework developed by Roze (2009) (see also Gervais et al. 2014 and Roze & Michod 2010) and explicitly considered chromosomes of size *L*, assuming that the S-locus was located at the center of the chromosome. Each generation, a random number of deleterious mutations was introduced along each chromosome, drawn from a Poisson distribution of parameter *U*, the haploid deleterious mutation rate. The position of each new deleterious mutation was drawn from a uniform continuous distribution, hence effectively allowing an infinite number of loci for deleterious mutations. At the reproduction stage, individuals produced recombinant gametes. The number of cross-overs occurring along the chromosome was drawn from a Poisson distribution of parameter *L* (genome map length in Morgans) and the position of each crossover from a uniform continuous distribution. All deleterious mutations were considered to have the same selection coefficient *s* and the same dominance coefficient *h*. The effect on gamete production *W i* of the deleterious mutations carried by individual *i* was supposed multiplicative, *i.e.* 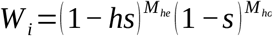, where *M _he_* and *M _ho_* are the number of mutations in the genome at heterozygous and homozygous states, respectively. We estimated inbreeding depression by 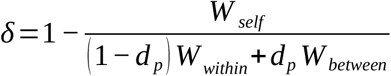 with *W_self_* the mean gamete production for an individual from selfing, *W_within_* the mean gamete production for an individual from a within-deme outcrossing and *W_between_* the mean gamete production for an individual from a between-demes outcrossing. Each mean gamete production is estimated from of a sample of 1,000 individuals, *i.e.* chromosome pairs, created independently of the main simulation process and with the appropriate mating rule.

#### Simulation procedure

In the model with constant inbreeding depression, simulations were first run until the effective number of SI alleles attained a drift-migration-mutation-selection equilibrium. We considered the equilibrium was reached when there was less than 2% variation of the average effective number of S-alleles over 200 generations. Self-compatible mutants were then introduced.

In the model with variable inbreeding depression, simulations were similarly run until the effective number of SI alleles reached equilibrium. Second, deleterious mutations were introduced and simulations were run until inbreeding depression reached mutation-drift-selection equilibrium. Third, mutations toward SC alleles were allowed. In both models, we determined the inbreeding depression threshold above which the SI system was maintained (less than 5 % of SC alleles at the end of the simulations) or below which it was lost because of the invasion of SC mutants (100% of SC alleles at the end of the simulation). In order to find the threshold value of interest, we explored the parameter space by modifying *δ* or *U* with a decreasing step size, increasing the value when SC mutants invaded and decreasing it when the SI system was maintained. Because of computation time constraints we stopped the exploration after a determined number of steps or if the SI system was not maintained despite convergence towards very high inbreeding depression (in the model where it was dynamic). We considered two conditions: 100% of SC allele despite *δ* ≥ 0.99 and more than 5% of SC allele despite *δ* ≥ 0.995. If one of these conditions was reached, we assumed that a fully functional SI system cannot be maintained.

## Results

In agreement with previous models (Charlesworth and Charlesworth 1979; Porcher and Lande 2005a; Gervais et al. 2014), our analytical model showed that the maintenance of SI was favored when the rate of self-pollen decreased and the number of SI alleles increased (Fig. 1, grey lines for *d_p_* = 0). Our model also showed that increasing the pollen dispersal rate *d_p_* rendered the maintenance of SI relatively easier. With *d_p_* = 10^*−*2^ the critical inbreeding depression needed to maintain SI was lower than for *d_p_* = 0 and this effect increased with an increasing number of SI alleles. Indeed, as the pollen dispersal rate increased, the amount of SI pollen received from the metapopulation and the amount of SC pollen exported to the metapopulation increased, which decreased fertilization opportunities for outcrossed SC pollen. In the extreme case of *d_p_* = 1, SI was always maintained, regardless of the level of inbreeding depression. Indeed, when *d_p_* = 1 outcrossing occurred only between demes such that all pollen from the focal deme was exported to the metapopulation and all outcrossed pollen in the focal population was SI pollen from the metapopulation. Globally, our analytical model showed that considering only the effect of pollen dispersal on pollen competition, dispersed pollen from other self-incompatible populations was expected to facilitate the local maintenance of SI. However, this model does not take into account the facts that 1) the SC mutation can actually occur anywhere in the metapopulation, not just the focal deme, 2) SC pollen from the focal deme can disperse and fertilize mates in other populations and 3) inbreeding depression and the number of SI alleles are not necessarily constant.

**Figure 1:**
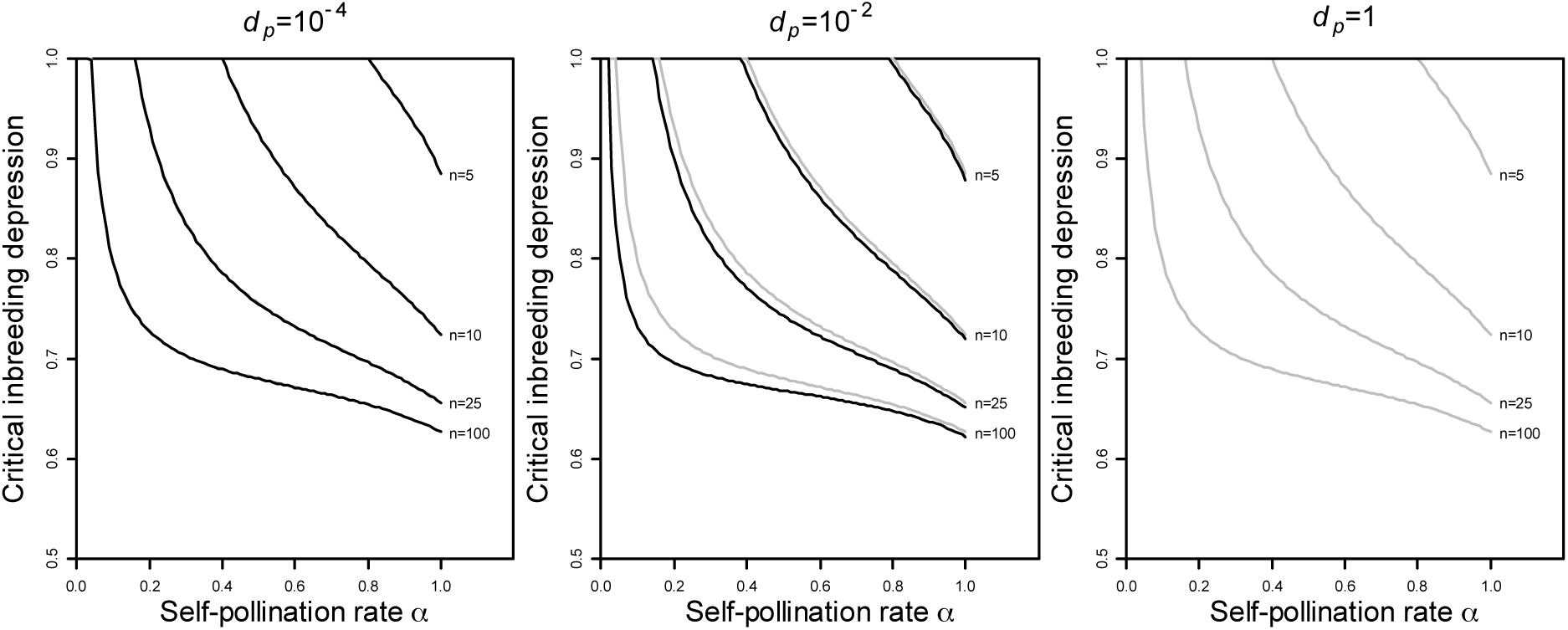
Analytical predictions of the critical inbreeding depression needed to maintain the SI system as a function of the self-pollination rate, for different numbers of SI alleles *n* in the focal deme and for *d_p_* = 10^*−*4^, *d_p_* = 10^*−*2^, *d_p_* = 1 (black curves) and for *d_p_* = 0 (grey curves). The SI system was maintained above the curves, while the frequency of SC mutants increased when rare below the curves. Note that for *d_p_* = 10^*−*4^, curves are superimposed with grey curves of *d_p_* = 0 and that for *d_p_* = 1, the SI system was always maintained (no black curve, only grey curves).

To explore these situations that are not tractable with an analytical model, we now show results obtained by stochastic simulations. In these simulations, the critical inbreeding depression required for SI maintenance consistently increased with increasing subdivision, *i.e.* decreasing pollen dispersal rate, for all self-pollination rates (Fig. 2). We have also investigated the effect of an increasing subdivision by rather increasing the number of demes and reducing their size and found the same result: the critical inbreeding depression required for SI maintenance increased with an increasing subdivision of the population. This effect is retained regardless of the self-pollen rate *α* (Fig. 3). In both cases, an increase in subdivision is associated with a non-monotonous variation of the global effective number of SI alleles (a decrease followed by an increase), but a monotonous decrease of the local effective number of SI alleles (Fig. 2b and 3b). These results are in line with Schierup (2000) and suggest that the monotonous increase we observed in the critical inbreeding depression required for SI maintenance is associated with the local effective number of SI alleles rather than the global effective number of SI alleles in the total population.

**Figure 2:**
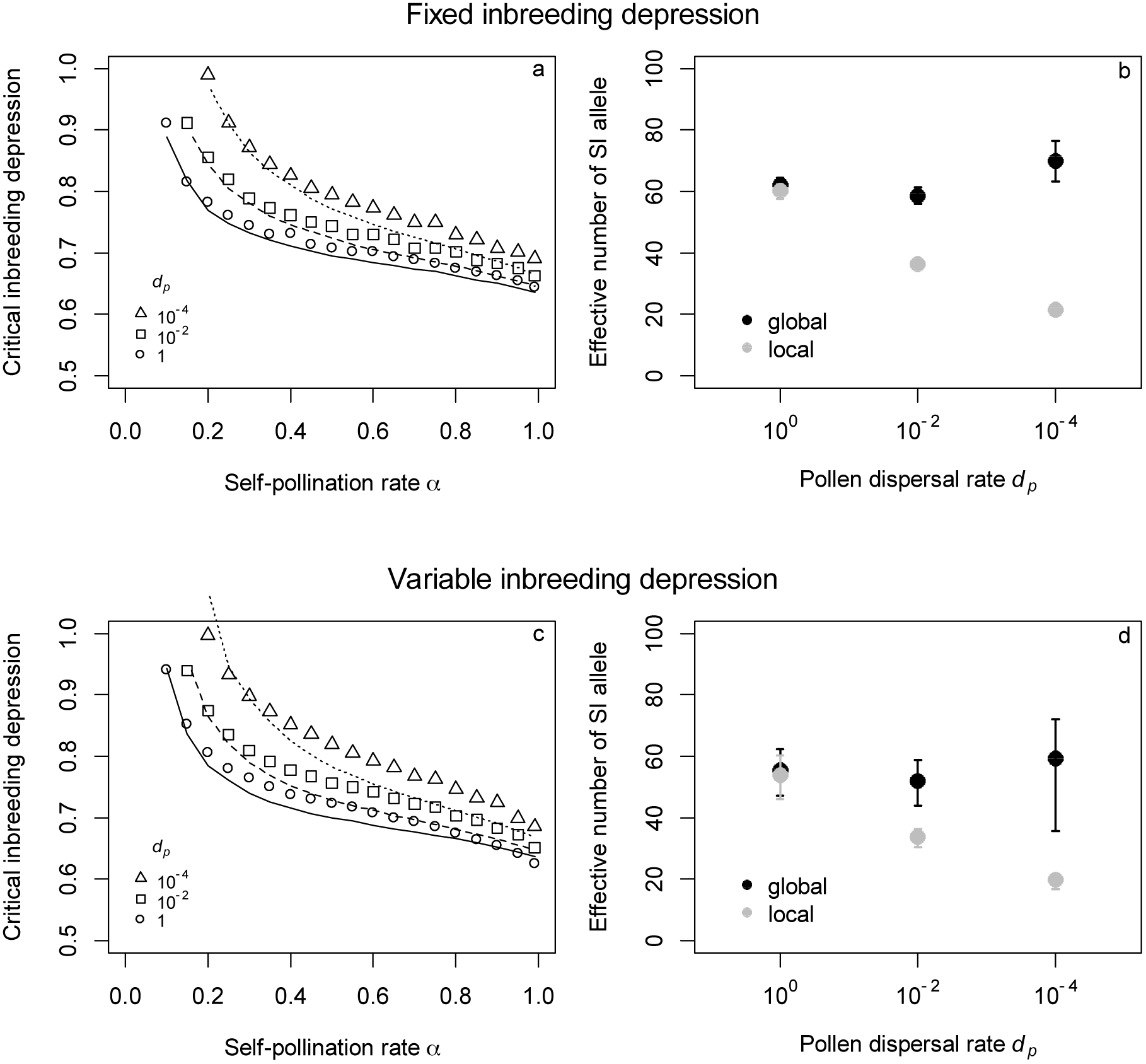
Effect of the rate of pollen dispersal on minimal inbreeding depression needed to maintain the SI system (a,c) and on the local and global effective numbers of SI alleles before introduction of SC mutants (b,d). (a,c) Symbols show simulation results. Each line shows analytical prediction for *d_p_* = 0 and for the median effective number of local allele obtained in the simulations (see panels b and d) for *d_p_* = 10^*−*4^ (dotted curve), *d_p_* = 10^*−*2^ (*dashed curve*) and *d_p_* = 1 (solid curve). (b,d) Bars show the distribution of 95% of the simulation results. (a,b) Constant inbreeding depression. (c,d) Variable inbreeding depression. Parameters (a,b,c,d) *N* = 10^5^ *, D* = 10, *S* = 1000,*m_SI_* = 10^*−*5^ *, m_SC_* = 10^*−* 4^. (c,d) *s* = 0.05,*h* = 0.2, *γ* = 0.5.

**Figure 3:**
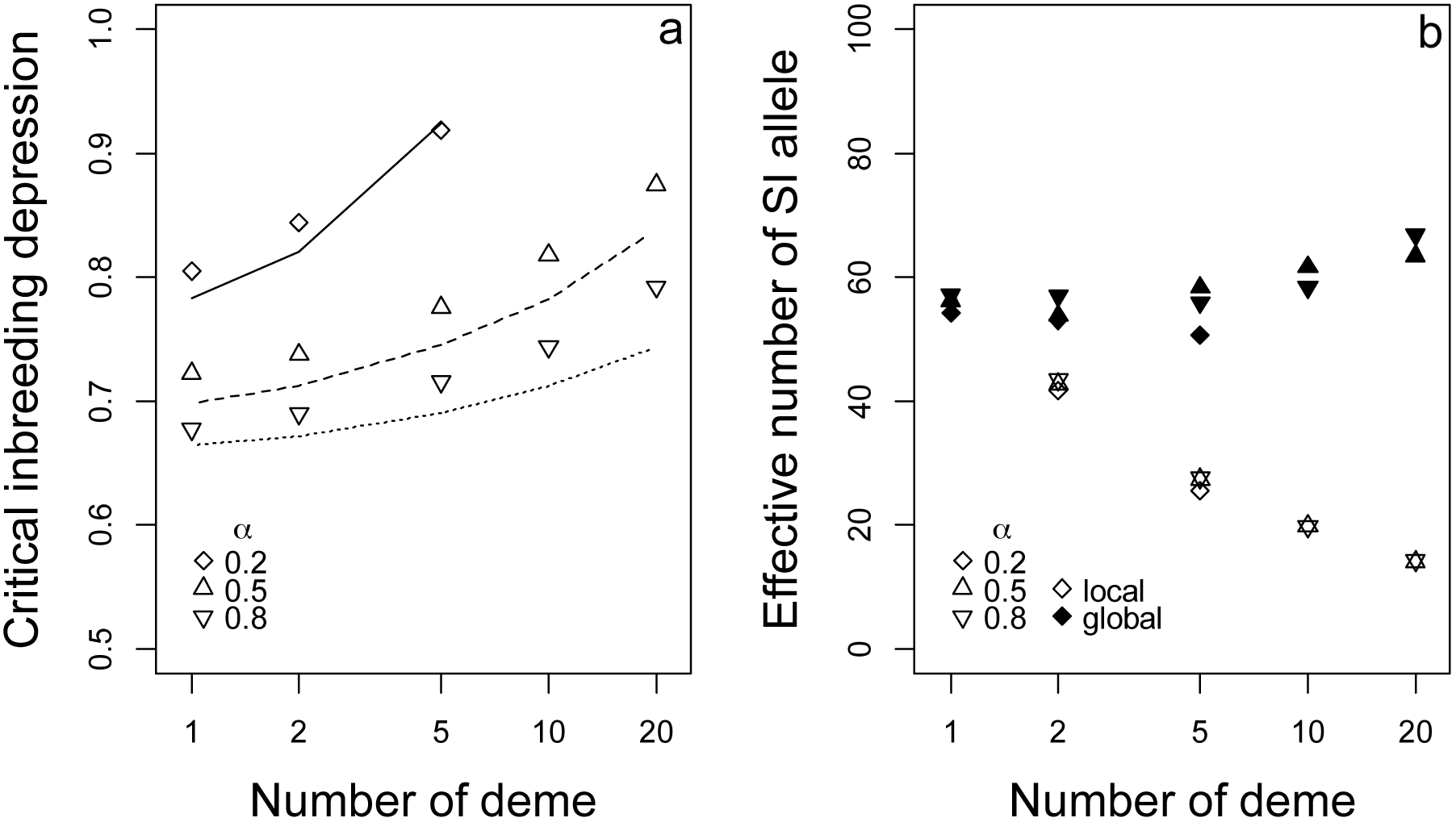
Effect of subdivision on critical inbreeding depression needed to maintain the SI system (a) and on the effective number of SI alleles before the introduction of SC mutants (b). (a) Symbols show simulation results. Each line shows analytical predictions for *d_p_* = 0 and for the effective number of local alleles obtained by simulation (see panel b) for *α* = 0.2 (dotted curve), *α* = 0.5 (dashed curve) and *α* = 0.8 (solid curve). (b) Effective local (open symbols) and global (closed symbols) number of SI alleles. Variable inbreeding depression. Parameters: *N* = 10^5^ *, d p* = 10^*−*4^ *, S* = 1000,*m_SI_* = 10^*−*5^ *, m_SC_* = 10^*−* 4^ *, s* = 0.05, *h* = 0.2, *γ* = 0.5.

In order to further test the hypothesis of the primary importance of the local number of SI alleles, we performed simulations with the same set of parameters except that we put a constraint on the number of possible SI alleles *S*. By reducing *S* we obtained similar local numbers of SI alleles despite the difference in subdivision (Fig. 4b, 4d and fig. 5b), allowing us to disentangle the effect of allelic richness from that of population subdivision itself. Our results showed that for a similar local number of SI alleles, the critical inbreeding depression needed to maintain SI was similar (Fig. 4a, 4c and fig. 5a). In other words, most of the effect of the spatial structure was captured by the local effective number of SI alleles. This hypothesis is supported by the comparison between simulations results and analytical predictions. Analytical predictions for a single isolated population (*d_p_* = 0) taking into account the number of SI alleles in the local populations were in good agreement with simulation results, for all dispersal rates (Fig. 2, 3). This again suggests that in our simulations, changes in the local number of S-alleles was the main driver of the change in inbreeding threshold of SI maintenance.

**Figure 4:**
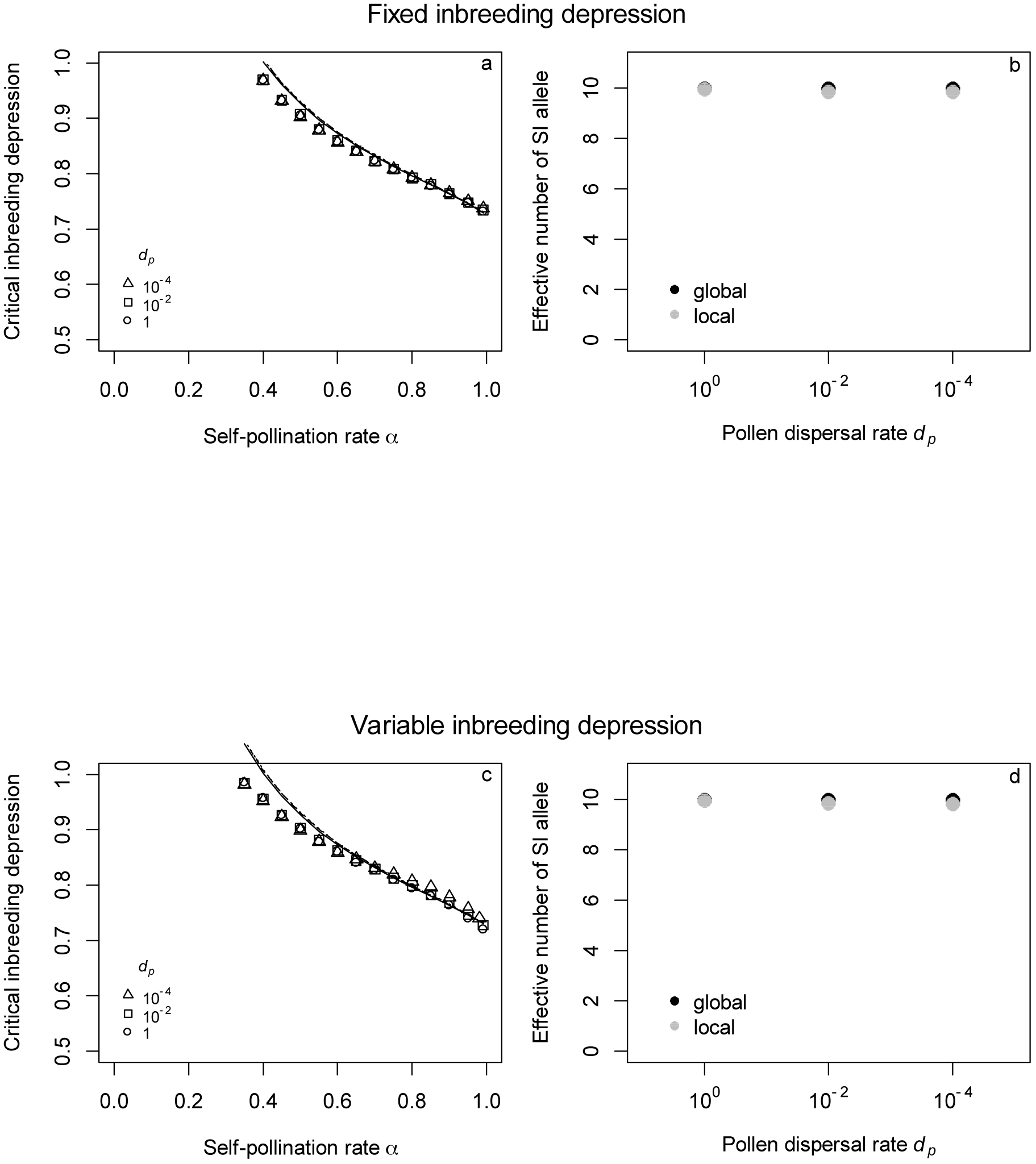
Effect of the rate of pollen dispersal on the minimal inbreeding depression needed to maintain the SI system (a,c) and on the local and global effective number of SI alleles before introduction of SC mutants (b,d) when the number of possible SI alleles *S* was constrained to low values (10). (a,c) Symbols show simulation results. Each line shows analytical predictions for *d_p_* = 0 and for the median effective number of local alleles obtained in the simulations (see panels b and d) for *d p* = 10^*−*4^ (dotted curve), *d p* = 10^*−*2^ (dashed curve) and *d_p_* = 1 (solid curve). (b,d) Bars showing the distribution of 95% of the simulations results were smaller than the symbols’ height and are therefore not represented. (a,b) Constant inbreeding depression (c,d) Variable inbreeding depression. Parameters (a,b,c,d) *N* = 10^5^, *D* = 10, *S* = 10,*m_SI_* = 10^*−*5^, *m_SC_* = 10^*−* 4^. (c,d) *s* = 0.05,*h* = 0.2, *γ* = 0.5.

**Figure 5:**
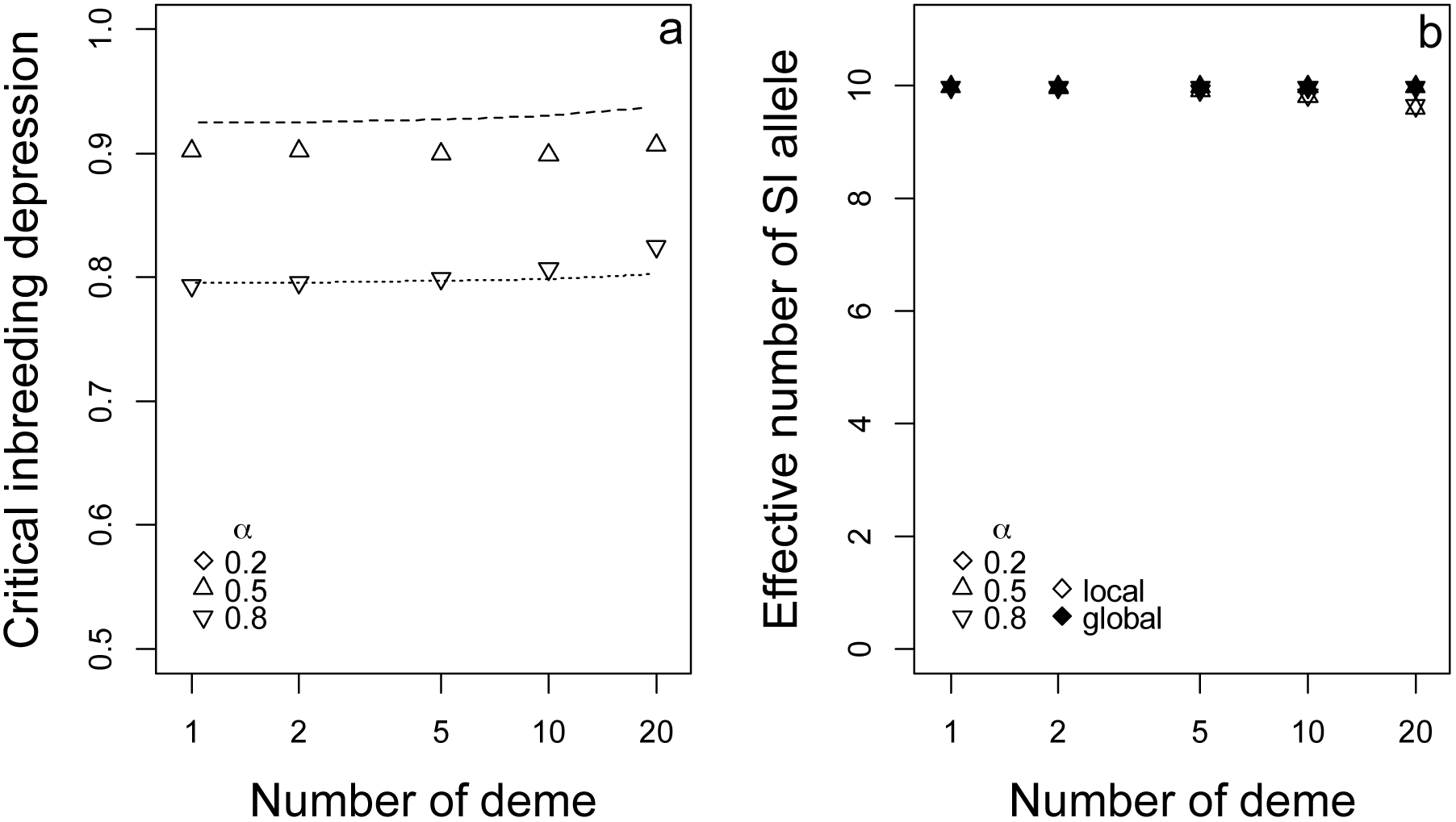
Effect of subdivision on the critical inbreeding depression needed to maintain the SI system (a) and on the effective number of SI alleles before introduction of SC mutants (b) when the number of possible SI alleles *S* was constrained to low values (10). (a) Symbols show simulation results. Each line shows analytical predictions for *d_p_* = 0 and for the effective number of local alleles obtained by simulation (see panel b) for *α* = 0.2 (no curve, the SI system was never maintained), *α* = 0.5 (dashed curve) and *α* = 0.8 (solid curve). (b) Effective local (open symbols) and global (closed symbols) numbers of SI alleles. Variable inbreeding depression. Parameters: *N* = 10^5^, *d p* = 10^−4^, *S* = 1000,*m_SI_* = 10^*−*5^, *mSC* = 10^*−* 4^, *s* = 0.05, *h* = 0.2, *γ* = 0.5.

Finally, in order to test whether spatial structure could affect the breakdown of SI through its effect on inbreeding depression, we compared the inbreeding depression threshold considering inbreeding depression either as a fixed parameter or as a variable depending on recurrent deleterious mutations. Overall, our results showed only very subtle quantitative differences between the two models. Specifically, the critical inbreeding depression needed to maintain SI was slightly higher in the model with variable inbreeding depression than in the model with fixed inbreeding depression (Fig. 2). This can be attributed to two phenomena: the effective number of local SI alleles was slightly lower with variable inbreeding depression (see Fig. 2b and 2d) and the partial purging of deleterious mutations (which should facilitate the invasion of SC mutants). Accordingly, under the parameters tested in our model, inbreeding depression was poorly affected by subdivision (Fig. 6). Globally, our results suggest that the effect of subdivision on inbreeding depression does not modify substantially the conditions for the maintenance of SI. This confirms that the most critical variable for the maintenance of SI is the local effective number of SI alleles.

**Figure 6:**
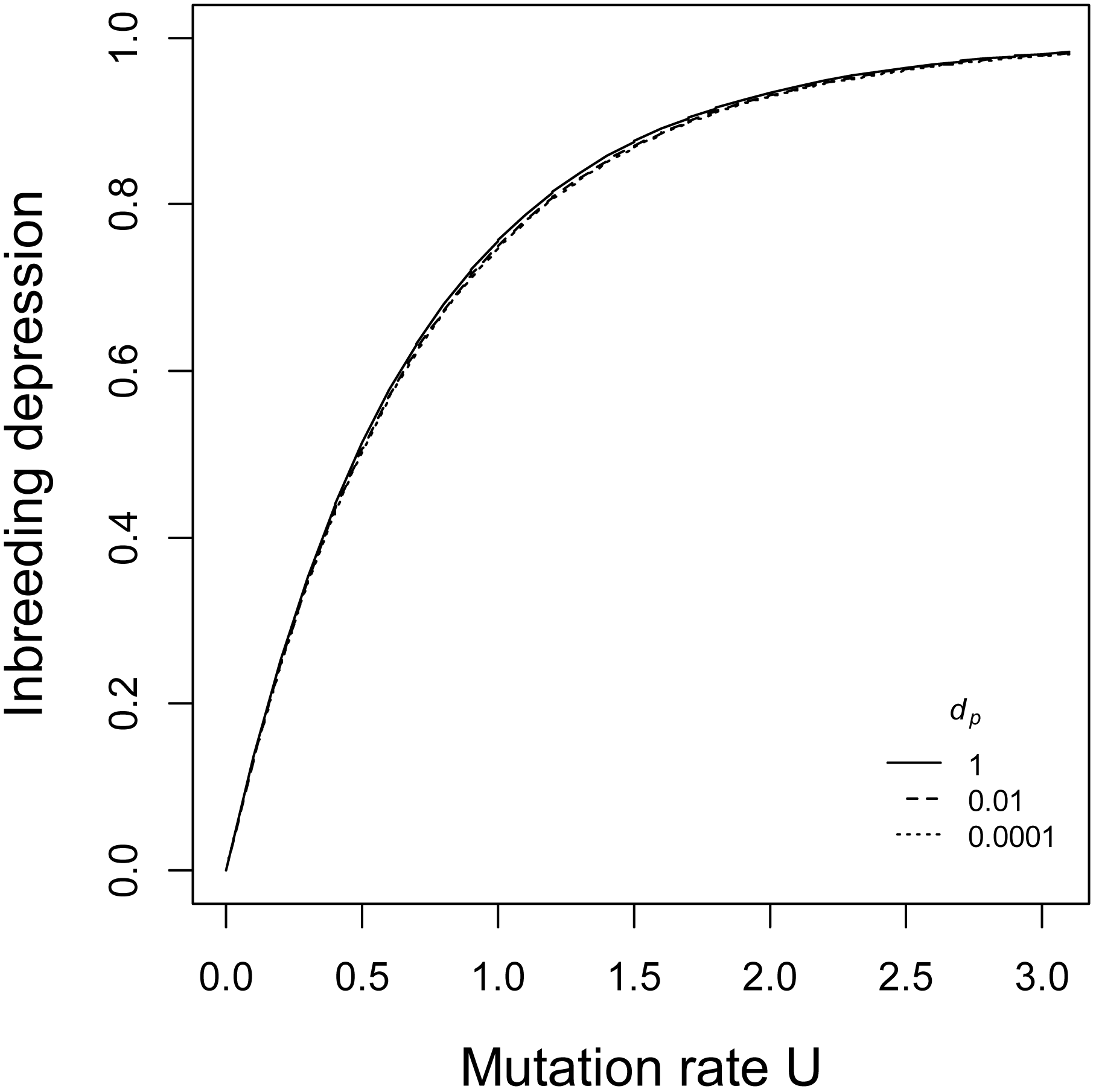
Inbreeding depression as a function of the deleterious mutation rate for different dispersal rates. Variable inbreeding depression. Parameters: *N* = 10^5^ *, D* = 10, *S* = 1000,*m_SI_* = 10^*−*5^ *, m_SC_* = 10^*−*4^ *, s* = 0.05, *h* = 0.2, *γ* = 0.5.

## Discussion

Our analytical and simulation results showed that population subdivision favors the breakdown of SI and that this effect mainly depends on the local diversity of SI alleles. At the deme scale, the effect of subdivision is therefore similar to the effect of a demographic bottleneck, with a loss of diversity of SI alleles. Bottlenecks are one of the main ecological factor proposed to explain transitions to self-compatibility (Reinartz and Les 1994; Porcher and Lande 2005a; Busch 2011). Such a bottleneck is expected at the range margin of species and in colonization processes as shown in *Arabidopsis lyrata* (Griffin and Willi 2014). A bottleneck also accompanied the transition to self-compatibility of *A. thaliana* (Durvasula et al. 2017) and *Capsella rubella* (Foxe et al. 2009). However, at the metapopulation scale, subdivision is expected to increase the effective population size (Wright 1943; Nei and Takahata 1993; Wang and Caballero 1999) and do not have the same effect than a bottleneck on SI-allele diversity (Schierup 1998). However, subdivision can have other effects that are not captured by our models. Empirical and theoretical analyses have shown that mating success of the female reproductive function of SI individual can been reduced in ecological conditions where compatible pollen is scarce (Busch and Schoen 2008; Leducq et al. 2010). Pollen limitation is indeed expected in spatially structured populations, especially at colonization fronts (Pannell 2015) or in poorly connected populations (Gascoigne et al. 2009). Self-compatibility is advantageous in conditions of pollen limitation because it can provide reproductive assurance (Busch and Delph 2012). While our models did not consider pollen limitation, we expect that it should further reduce the conditions for SI maintenance (Porcher and Lande 2005a), hence reinforcing the effect of population structure we documented, or promote stable mixed mating system (Porcher and Lande 2005b). In addition, the comparison between our analytical predictions and simulations results show that under the assumption of our analytical model the effect of pollen flow on pollen competition can greatly improve SI maintenance independently of an increase in the number of SI alleles. The assumption of our analytical model of neglecting pollen exported from the focal deme can be compared to pollen exchange between a marginal and a core population that SC alleles cannot invade, a situation that our individual-based models did not capture and that it might be interesting to investigate.

We did not see a strong qualitative effect of considering inbreeding depression as a fixed parameter or as a dynamic variable. This suggests that the purging effect is minor, and only slightly affected by population subdivision. This result is consistent with the literature because we considered weak effect recessive deleterious mutations for which purging by selfing is limited (Gervais et al. 2014). Our model assumed that inbreeding depression was caused by recessive deleterious mutations (Charlesworth and Willis 2009), and a possible extension of our model would be to implement other types of mutations, such as highly deleterious or lethal recessive mutations. For these mutations, we expect a more intense purging effect (Gervais et al. 2014; Roze 2015; García-Dorado 2017), so we predict that the introduction of such mutations would lead to even more restricted conditions for the maintenance of SI.

Our results have important implications for the interpretation of empirical data on the loss of SI in natural populations. In fact, population subdivision is one the most common features of natural populations, and the level of subdivision can vary through time for a given species. As explained above, the effect of subdivision on SI maintenance is akin to a genetic bottleneck, which has been considered one of the main ecological factors to explain transitions to self-compatibility (Reinartz and Les 1994; Porcher and Lande 2005a; Busch 2011). Such bottlenecks are expected at the range margin of species and in colonization processes as shown in *Arabidopsis lyrata* (Griffin and Willi 2014). A bottleneck also accompanied the transition to self-compatibility of *A. thaliana* (Durvasula et al. 2017) and *Capsella rubella* (Foxe et al. 2009). Furthermore, shifts of the climatic conditions can modify the distribution range as well as alter the connectivity among demes (Thomas et al. 2004). Similarly, the loss of an important pollinator may modify the conditions of pollen transport (Aguilar et al. 2006). These effects were not previously integrated in models for the maintenance of SI, and our results predict that these ecological factors should strongly and directly affect the maintenance of SI. Several lines of evidence suggest that the breakdown of SI has indeed occurred in spatially structured populations in several plant species. For instance, North American populations of *A. lyrata* show multiple independent breakdowns of SI (Foxe et al. 2010) and distinct SC mutations have been found across Leavenworthia (Busch et al. 2011) and Solanum (Markova et al. 2016) populations. In the selfer *A. thaliana*, as many as three different causal segregating SC mutations have been identified, possibly associated with distinct ancient glacial refugia (Shimizu et al. 2008; Durvasula et al. 2017; Tsuchimatsu et al. 2017). In light of our results, we propose the interpretation that these observations correspond to cases where subdivision has increased species-wide because of changes of ecological factors, bringing the conditions to SC invasion all over the range, hence allowing invasion by multiple SC mutants.

Our results also have important implications in the context of how new SI alleles can arise through allelic diversification (Uyenoyama et al. 2001). Because the male and female components of SI are typically encoded by distinct (but tightly linked) genes (Takayama and Isogai 2005), allelic diversification has been proposed to arise through the transient segregation of mutants that bear mutations modifying recognition specificity of one of the two genes only, compromising the male-female recognition and resulting in SC alleles. The conditions under which this initial SC mutation can stably segregate without being either eliminated or fixed are crucial to predict whether it can be hit by a mutation on the second gene creating a novel recognition specificity, resulting in allelic diversification (Uyenoyama et al. 2001; Gervais et al. 2011). As shown by Gervais et al. (2011), the dynamics of diversification of SI alleles decreased as the number of SI alleles itself increased, so we expect that population subdivision, by keeping the number of SI alleles at a low level may actually increase the rate at which new SI alleles arise. Moreover, Gervais et al. 2011 also predicted that in a large portion of the parameter range the transient SC allele was expected to replace its ancestral copy, resulting in allelic turnover rather than diversification *per se*. It is possible that in a metapopulation, the replacement phenomenon may take place independently in the different demes if they are sufficiently isolated, resulting in the origin of new alleles in different parts of the range. It will now be interesting to study whether this process indeed allows for the increase in diversity of SI alleles in a wider range of parameters than in a single isolated population.

To conclude, we showed that spatial structure is an important factor affecting the conditions for the maintenance of SI. We demonstrated that, under the parameters tested in our study, it is the effect of population subdivision on the number of SI alleles locally maintained in the demes that determines the fate of SC mutants when they arise rather than a change in the dynamics of inbreeding depression. Our results have implications for the interpretation of the scenarios by which plant mating systems can shift in natural populations and may change our understanding of the factors affecting the process of allelic diversification.

Author contributions
TB and SB designed the models. TB conducted the simulations and analyzed the models. SB and VC supervised the work.

## Acknowledgements

This work was funded by the European Research Council (NOVEL project, grant #648321). The authors also thank the Région Hauts-de-France, and the Ministére de l’Enseignement Supèrieur et de la Recherche (CPER Climibio), and the European Fund for Regional Economic Development for their financial support.

